## Supplementary figures and images for "Retrograde And Anterograde Transport Of LAT-Vesicles During The Immunological Synapse Formation: Defining The Finely-Tuned Mechanism"

### Supplementary Figures.pdf

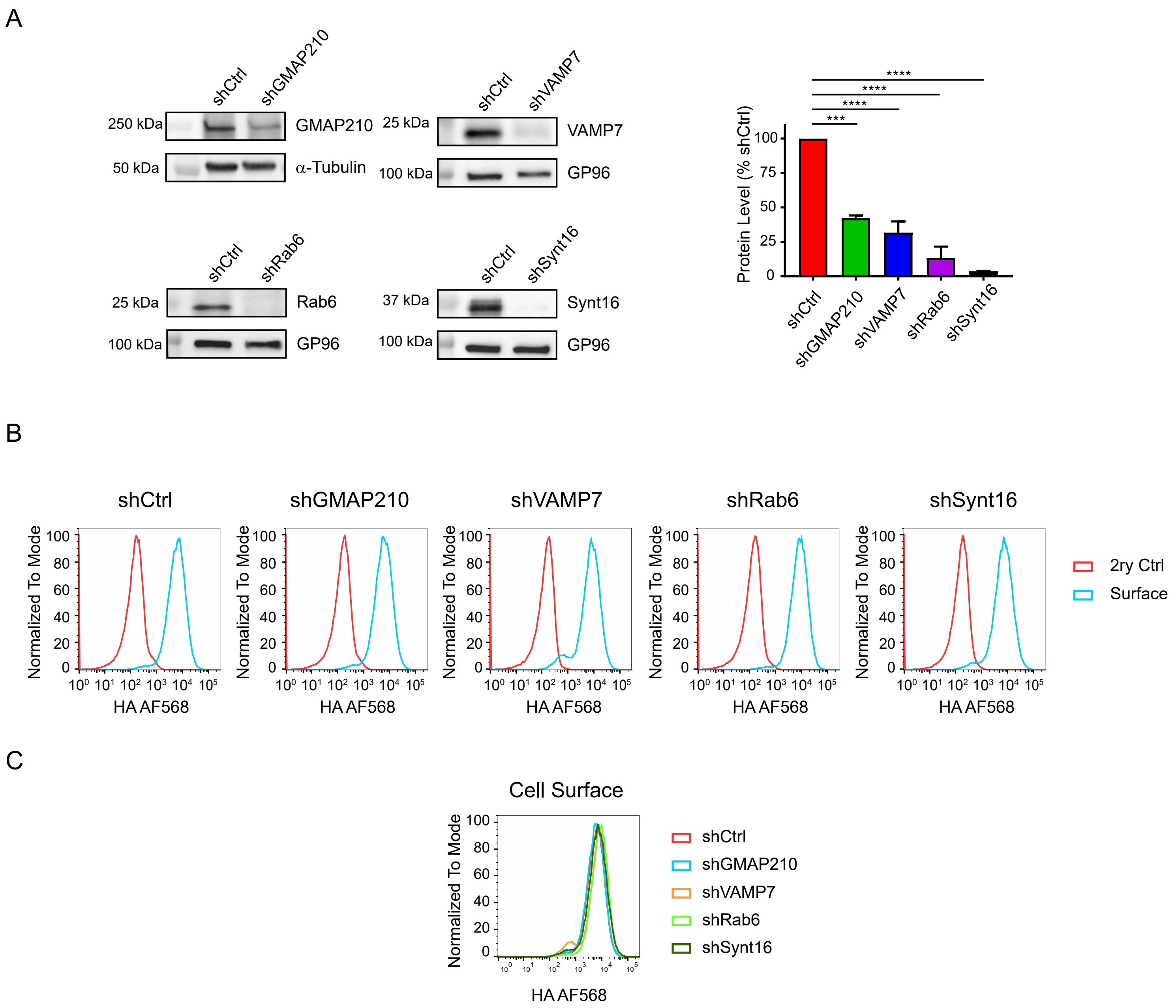

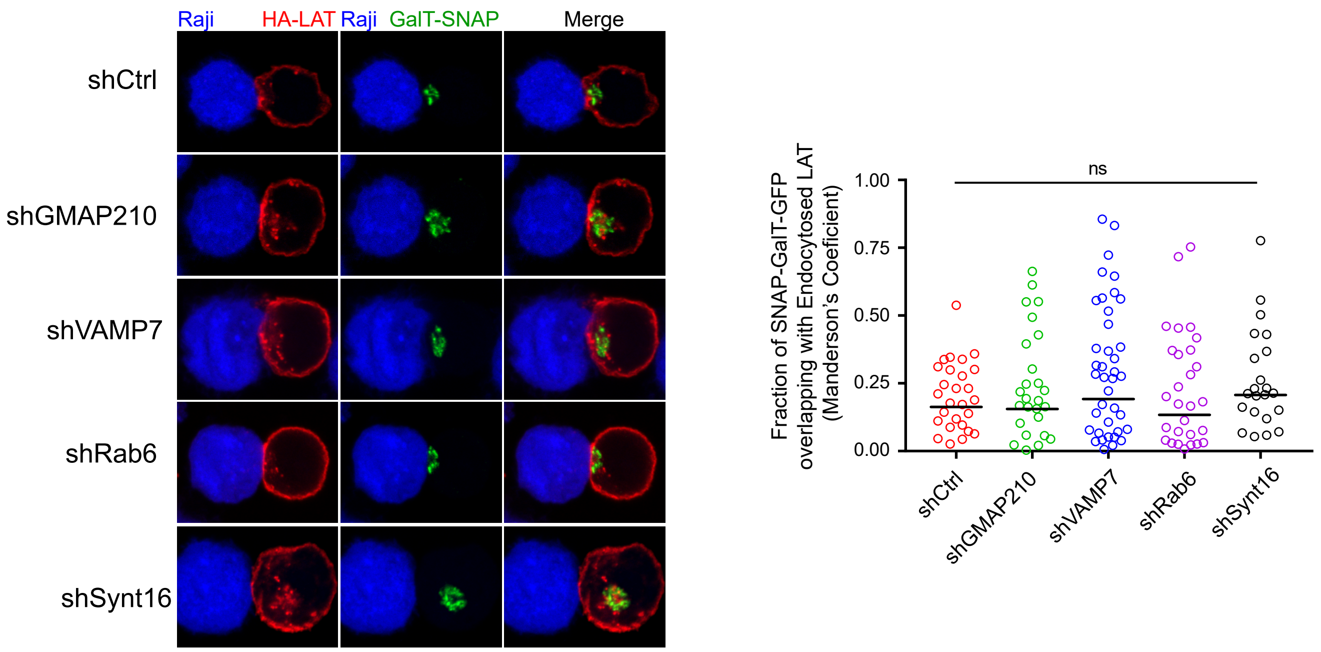

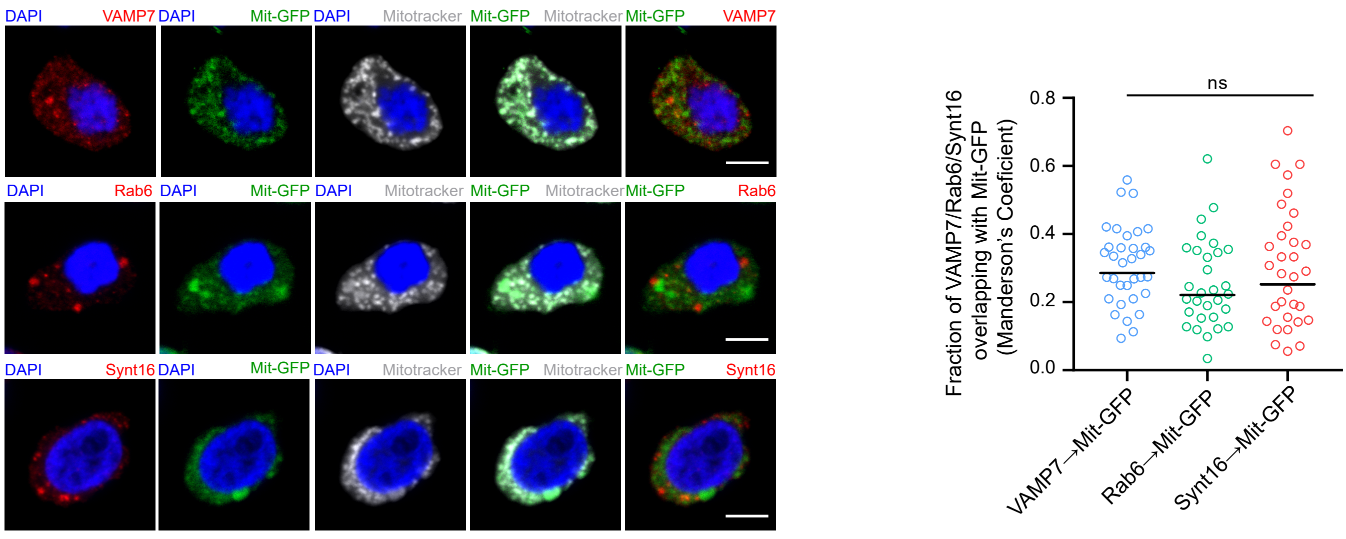
